# Sequence neighborhoods enable reliable prediction of pathogenic mutations in cancer genomes

**DOI:** 10.1101/2021.02.09.430460

**Authors:** Shayantan Banerjee, Karthik Raman, Balaraman Ravindran

**Affiliations:** Robert Bosch Centre for Data Science and Artificial Intelligence (RBCDSAI), Indian Institute of Technology (IIT) Madras, Chennai - 600 036; Initiative for Biological Systems Engineering, IIT Madras, Chennai - 600 036; Bhupat and Jyoti Mehta School of Biosciences, Department of Biotechnology, IIT Madras, Chennai - 600 036; Department of Computer Science and Engineering, IIT Madras, Chennai - 600 036

## Abstract

Identifying cancer-causing mutations from sequenced cancer genomes hold much promise for targeted therapy and precision medicine. “Driver” mutations are primarily responsible for cancer progression, while “passengers” are functionally neutral. Although several computational approaches have been developed for distinguishing between driver and passenger mutations, very few have concentrated on utilizing the raw nucleotide sequences surrounding a particular mutation as potential features for building predictive models. Using experimentally validated cancer mutation data in this study, we explored various string-based feature representation techniques to incorporate information on the neighborhood bases immediately 5’ and 3’ from each mutated position. Density estimation methods showed significant distributional differences between the neighborhood bases surrounding driver and passenger mutations. Binary classification models derived using repeated cross-validation experiments gave comparable performances across all window sizes. Integrating sequence features derived from raw nucleotide sequences with other genomic, structural and evolutionary features resulted in the development of a pan-cancer mutation effect prediction tool, NBDriver, which was highly efficient in identifying pathogenic variants from five independent validation datasets. An ensemble predictor obtained by combining the predictions from NBDriver with two other commonly used driver prediction tools (CONDEL and Mutation Taster) outperformed existing pan-cancer models in prioritizing a literature-curated list of driver and passenger mutations. Using the list of true positive mutation predictions derived from NBDriver, we identified a list of 138 known driver genes with functional evidence from various sources. Overall, our study underscores the efficacy of utilizing raw nucleotide sequences as features to distinguish between driver and passenger mutations from sequenced cancer genomes.

## Introduction

Cancer is caused due to the accumulation of somatic mutations during an individual’s lifetime [1]. These mutations arise due to both endogenous factors such as errors during DNA replication, or exogenous factors such as substantial exposure to mutagens such as tobacco smoking, UV light, and radon gas. [2]–[4]. These somatic mutations can be of different types, ranging from single-nucleotide variants (SNVs), to insertions and deletions of a few nucleotides, copy-number aberrations (CNAs), and large-scale rearrangements known as structural variants (SVs) [5]. With the advent of high-throughput sequencing, the identification of somatic mutations from sequenced cancer genomes has become easier. International cancer genomics projects have resulted in the development of large mutational databases such as the Catalogue Of Somatic Mutations In Cancer (COSMIC) [6], the International Cancer Genome Consortium (ICGC) [7], and The Cancer Genome Atlas (TCGA) [8]. Several open-access resources to analyze and visualize large cancer genomics datasets, such as the cBio Cancer Genomics Portal [9] and the Database of Curated Mutations in cancer (DoCM) [10], have also been developed. These resources aggregate functionally relevant cancer variants from different studies and help researchers gain easy access to expert-curated lists of pathogenic somatic variants.

However, not all somatic mutations present in the cancer genome are *equally* responsible for developing the disease. A small fraction of somatic variants known as “driver mutations” provide a growth advantage and are positively selected for, during cancer cell development [1]. On the other hand, “passenger mutations” provide no growth advantage and do not contribute to cancer progression [1]. Identifying the complete set of cancer-causing genes that harbor driver mutations, also known as driver genes, holds much promise for precision medicine, where a specific therapeutic intervention is tailored towards a patient’s mutational profile [11].

Distinguishing between driver and passenger mutations from sequenced cancer genomes is a non-trivial task. Doing so solely based on the substitution type (A->T, G->C, etc.) is very difficult. Hence, several computational methods that utilize several other factors to identify driver mutations have been developed over the years. Recurrence-based driver prioritization tools such as MutSigCV [12] and MuSiC [13] for single-nucleotide variants, and GISTIC2 [14] for copy number aberrations, have been developed to identify variants that occur more than what is expected by chance, otherwise known as the “background mutation rate”. Other methods such as SIFT [15], PROVEAN [16], PolyPhen-2 [17], CHASM [18], and FATHMM [19] are based on predicting the functional impact of mutations on the protein encoded by the gene. Expert-curated databases such as the OncoKB database [20] contain information regarding the functional impact of over 3000 cancer-causing alterations belonging to over 400 genes. Pathway analysis based tools such as NetBox [21] and HotNet [22] work by identifying mutations affecting large scale gene regulatory or protein–protein interaction networks. Machine learning-based methods have also been recently developed to predict deleterious missense mutations [23]–[28].

Genome instability, demonstrated by a higher than average rate of substitution, insertion, and deletion of one or more nucleotides, is a hallmark of most cancer cells. There is a considerable variation in the rates of SNPs across the human genome. Sequence context plays a significant role in the variability of the substitutions rate as explained by the CpG dinucleotides, which exhibit an elevated C->T substitution rate by almost 15 folds relative to the average rate observed in mammals [29]. Mutational hotspots such as the CpG dinucleotides in breast and colorectal cancer [30] and TpC dinucleotides in lung cancer, melanoma, and ovarian cancer [31] are some examples of “signatures” that promote mutagenesis. There have been several efforts to utilize the sequence context to measure the human genome’s substitution rates. Aggarwala *et al*. [32] used the local sequence context of SNPs to explain the observed variability in substitution rates. Zhao *et al*. [33] studied the neighboring nucleotide biases and their effect on the mutational and evolutionary processes for over two million SNPs.

Recent studies have identified specific signatures or patterns of mutations in different cancer types that shed light on the underlying mechanisms responsible for cancer progression [34], [35]. Alexandrov *et al*. [34] identified 21 distinct mutational signatures in human cancers by considering the substitution class and the sequence context immediately to the 3’ and 5’ of the mutated base. Several studies have demonstrated that certain factors such as tobacco smoking, UV light, or the inactivation of tumor suppressor genes involved in DNA repair can result in the development of mutational hotspots [31], [34], [36]. There have been two recently published studies that have tackled this problem, to the best of our knowledge. Deitlein *et al*. [61] hypothesized that driver mutations occur more frequently in “unusual” nucleotide positions than passenger mutations and built probabilistic models to identify driver genes that had mutations in those “unusual” contexts. Agajanian *et al*. [37] integrated classical machine learning and deep learning approaches to model raw nucleotide sequences to differentiate between driver and passenger mutations.

In this study, our overall aim is to build models utilizing machine learning and natural language processing techniques to differentiate between driver and passenger mutations solely based on the raw nucleotide context. Using missense mutation data with experimentally validated functional impacts compiled from various studies, we show that the underlying probability distributions of driver and passenger mutations’ neighborhoods are significantly different from one another. We extracted features from the neighborhood nucleotide sequences and built robust binary classification models to distinguish between the two classes of mutations. We achieved good classification performances during our repeated cross-validation experiments and against an independent hold-out set of literature curated mutations. Integrating neighborhood features with other features such as protein physicochemical properties and evolutionary conservation scores significantly improved our algorithm’s overall predictive power in identifying pathogenic variants from five separate independent test sets, and had comparable performances with some of the existing state-of-the-art mutation effect prediction tools. Overall, this study establishes that we can leverage efficient feature representation of the neighborhood sequences of cancer-causing mutations to differentiate between a known driver and passenger mutations with sufficient discriminative power.

## Methods

### Mutation Datasets for Building and Evaluating the Models

Our training data consisted of the list of missense mutations whose effects were determined from experimental assays and were compiled in the study conducted by Brown *et al*. [37]. In this study, missense mutations from 58 genes that were pan-cancer-based were combined from five different datasets [38], [75]–[79] (Supplementary Table 1). These mutations were presented as amino acid substitutions based on their protein coordinates (e.g., F595L, L597Q, etc.). Since we were interested in studying the effects of neighboring DNA nucleotide sequences, we mapped them to their corresponding genomic coordinates (gDNA) for further analysis. We used the publicly available TransVar web-interface [80] for this purpose. The final training set was made up of 5265 single nucleotide variants (4131 passengers and 1134 drivers).

For external validation, we collected somatic mutation data from five different sources. First, we considered a literature-curated list of 140 passengers and 849 driver mutations categorized based on functional evidence published by Martelotto *et al*. [38] as part of the benchmarking study to rank various mutation effect prediction algorithms.

Second, we used a subset of mutations published by the recently released Cancer Mutation Census. The Cancer Mutation Census (CMC) [6] is a database that integrates all coding somatic mutation data from the COSMIC database to prioritize variants driving different cancer forms. It contains functional evidence obtained using both manual curation and computational predictions from multiple sources. For our validation experiments, we chose only single nucleotide variants classified as missense and derived from the CGC-classified list of tumor suppressor genes and oncogenes. Based on the database’s various evidence criteria, we considered only mutations categorized as tier 1, 2, and 3 for our study. From this list, we further removed all overlapping mutations with our training set and derived a final set of 277 mutations for further analysis.

The Catalog of Validated Oncogenic Mutations from the Cancer Genome Interpreter [35] database contains a high confidence list of pathogenic alterations compiled from several sources such as the DoCM [10], ClinVar [81], OncoKB [20], and the Cancer Biomarkers Database [35]. We extracted only missense somatic mutations flagged as “cancer” for our validation experiments. After removing all overlapping mutations with our training set, we obtained a final list of 1628 driver mutations. This constituted our third validation set.

The fourth validation dataset consisted of the list of top 50 hotspot mutations reported in the comprehensive study done by Rheinbay *et al*. [44]. In this study, mutation data was accumulated from the Pan-Cancer Analysis of Whole Genomes (PCAWG) consortium and involved analyzing more than 2700 cancer genomes derived from more than 2500 patients. A total of 33 coding missense mutations from five well-known cancer genes: TP53, PIK3CA, NRAS, KRAS, IDH1, were extracted from this study.

Mao *et al*. [27] published mutation datasets to judge the performance of the driver prediction tool (CanDrA) in predicting rare driver mutations. They were constructed using the following criteria:

1. GBM and OVC mutations reported in the COSMIC database only once.
2. The reported mutations had no other mutations within 3bp of their position and were not part of either the training or test datasets for building the machine learning model (CanDrA).

We used the same datasets to judge our model’s ability to predict rare driver mutations based solely on the neighborhood sequences. After removing all overlapping mutations with the training set, we obtained 34 GBM mutations and 38 OVC mutations. A summary of all the mutational datasets used in our study is available in Table 1. Besides, all our predictions are derived using the forward strand and were based on the GRCh37 (ENSEMBL release 87) build of the human genome.

**Table 1:** Summary of datasets used in this study

| Type | Study/<br>Database Name | Description | Sample size |
| --- | --- | --- | --- |
| Training | Brown *et al.* | Missense mutations from 58 cancer genes generated from experimental assays | 5265 mutations<br>(Driver: 1134<br>Passenger: 4131) |
| Validation | Martelotto *et al.* | A Literature curated list of mutations from 15 cancer genes used to benchmark 15 mutation-effect prediction algorithms | 989 mutations<br>(Driver: 849<br>Passenger: 140) |
| Validation | Catalog of Validated Oncogenic Mutations | High confidence pathogenic missense variants compiled from several sources | 1628 driver mutations |
| Validation | Rheinbay *et al.* | Recurrent single point driver mutations in the coding region compiled from the Pan-Cancer Analysis of Whole Genomes Consortium | 33 driver mutations |
| Validation | Mao *et al.* | Rare driver mutations from GBM and OVC cancer types | GBM: 34 driver mutations<br>OVC: 38 driver mutations |
| Validation | Cancer Mutation Census (COSMIC v92) | COSMIC mutation data categorized into different functional classes both through manual curation and computational predictions | 277 driver mutations |

### Feature Extraction

#### Sequence-Based Features

We used the raw nucleotide sequences surrounding a mutation as features for our analysis. Each unique mutation was represented as a triplet (Chromosome, Position, Type) where “Type” refers to one of the 12 types of point substitution (A>T, A>G, A>C T>A, T<G, T>C G>A, G>C, G>T, C>T, C>A, C>G). We then extracted the surrounding raw nucleotide sequences from the reference genome for a given mutation position using the bedtools getfasta command. The “window size” for a particular mutation captures the number of nucleotides upstream and downstream from the mutated position. Hence, considering all possible window sizes between 1 and 10, including the wild-type nucleotide at the mutated position, we obtained nucleotide strings of length 3, 5, 7, 9, 11, 13, 15, 17, 19, and 21, respectively. We also considered the chromosome number and the type of point substitution as features for our analysis. Now, for particular window size, to map the nucleotide strings to a numerical format, we used the following two widely used feature transformation approaches (Figure 1):

1. **One-hot encoding**: Each neighboring nucleotide was represented as a binary vector of size 4 containing all zero values except the nucleotide index, which was marked as 1. Thus “A” was encoded as [1,0,0,0], “G” as [0,1,0,0] and so on. This particular feature representation resulted in a feature space of size *8n* + 2, where *n* = 1,2,3 … 10 represents the window sizes. We used the pandas `get_dummies()` to perform this task.
2. **Overlapping *k*-mers**: In this type of feature representation, the neighboring nucleotide string sequences for a given window size were represented as overlapping *k-*mers of lengths 2,3 and 4. For instance, an arbitrary sequence of window size 3 {ATT**T**GGA}, where ‘**T**, is the wild type base at the mutated position, can be decomposed into overlapping *k-mers* of size 2 {AT, TT, T**T**, **T**G, GG, GA}, 3 {ATT, TT**T**, T**T**G, **T**GG, GGA} and 4 {ATT**T**, TT**T**G, T**T**GG, TGGA} respectively.

**Figure 1:**
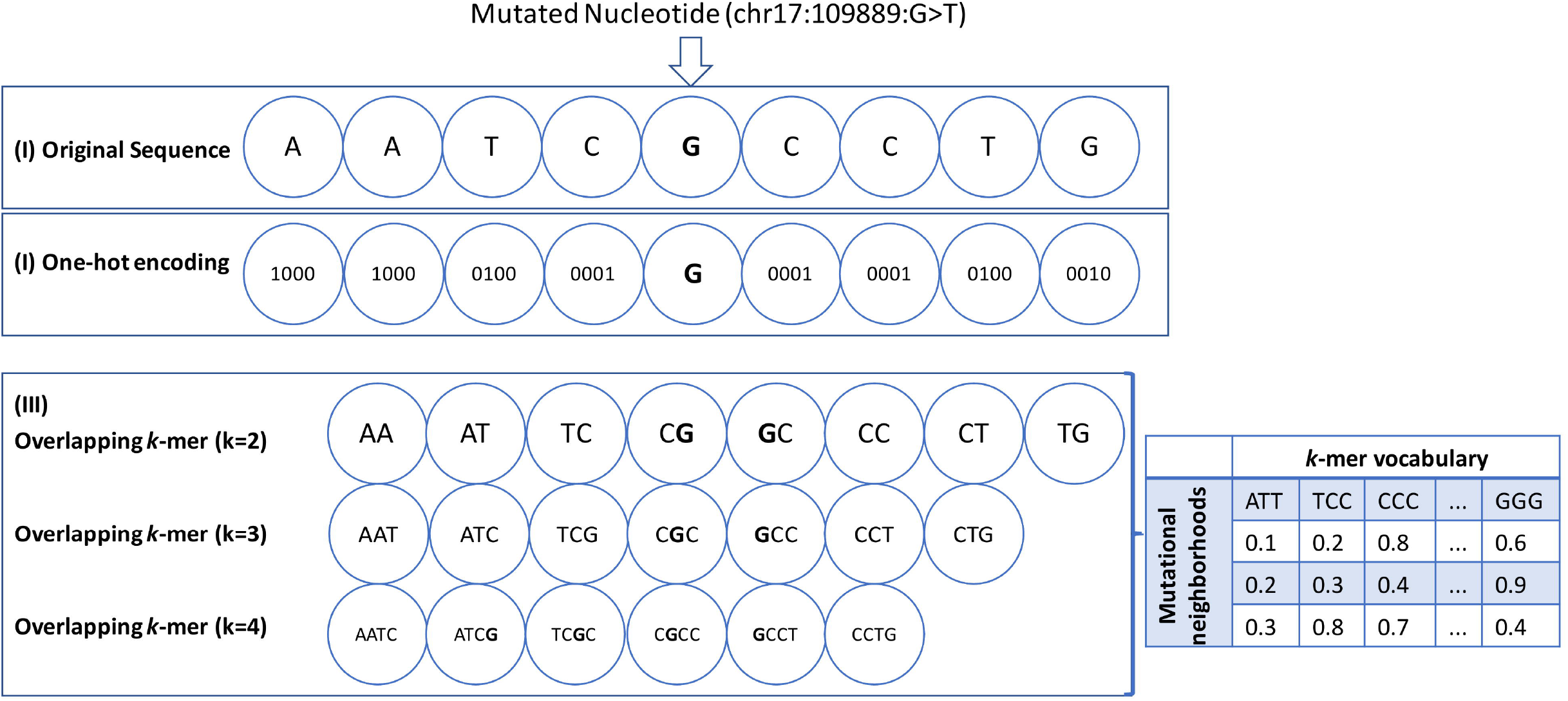
A diagram representing the features derived from the neighborhood nucleotide sequences of the point mutations for an arbitrary window size of 4 is shown here. The mutated position is represented as a triplet (Chromosome: Position: Substitution Type). **(I)** The original sequence is represented here with the mutated nucleotide (ch17:109889:G>T) in bold. **(II)** One-hot encoding was used to derive the 4-bit binary one-hot encoded vector for each nucleotide. **(III)** Overlapping *k*-mers of sizes 2,3 and 4 have been represented here. In this case, the neighborhood features also include the wildtype nucleotide at the mutated position. The overlapping *k*-mers were encoded into a numerical format using the countvectorizer and the TFIDF vectorizer and the resulting word matrix was derived. The samples (or individual neighborhoods) are represented as rows and the *k*-mers are represented as columns. For both types of feature representation, the chromosome number and the substitution type (A>T, G>C etc) were included as additional features.

To map these overlapping *k*-mers to a numerical format, we applied two commonly used encoding techniques known as CountVectorizer and TfidfVectorizer. The CountVectorizer returns a vector encoding whose length is equal to that of the vocabulary (total number of unique *k*-mers in the data set) and contains an integer count for the number of times a given *k*-mer has appeared in our dataset.

A Term Frequency – Inverse Document Frequency (TF-IDF) vectorizer assigns scores to each *k*-mer based on i) how often the given *k*-mer appears in the dataset and ii) how much information the given *k*-mer provides, i.e., whether it is common or rare in our dataset. Mathematically, for a given term *i* present in a document*j*, the TF-IDF score *tf_i,j_* is given by

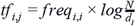

where *freq_i,j_* is the number of occurrences of *i* in *j*, *d_i_* is the number of documents containing *i*, and *N* is the total number of documents. These techniques were implemented in Python using the *feature_extraction* module from scikit-learn. The final processed training set used to build the machine learning models was represented as a matrix of size *mn*, where *m* is the total number of coding point mutations and *n* is the size of the vocabulary. The matrix entries were the TF-IDF or the CountVectorizer scores. The number of one-hot encoded features, *k*-mers, and the size of the vocabulary possible for each window size is shown in Table 2.

**Table 2:** Number of one-hot encoded features and possible *k*-mers for a given window size. The size of the vocabulary (or **N**) is given in brackets

| Window Size | Number of one-hot encoded features | Number of $k$-mers possible for a given $k$-mer size | | |
| --- | --- | --- | --- | --- |
| | | $k=2$<br>($N=16$) | $k=3$<br>($N=64$) | $k=4$<br>($N=256$) |
| w= 1 | 8 | 2 | 1 | 0 |
| w= 2 | 16 | 4 | 3 | 2 |
| w= 3 | 24 | 6 | 5 | 4 |
| w= 4 | 32 | 8 | 7 | 6 |
| w= 5 | 40 | 10 | 9 | 8 |
| w= 6 | 48 | 12 | 11 | 10 |
| w= 7 | 56 | 14 | 13 | 12 |
| w= 8 | 64 | 16 | 15 | 14 |
| w= 9 | 72 | 18 | 17 | 16 |
| w= 10 | 80 | 20 | 19 | 18 |

#### Descriptive Genomic Features

In addition to the neighborhood features, a set of 27 features (Supplementary Table 2) previously used to train the cancer-specific missense mutation annotation tool, CanDrA [27], were extracted from the following three data portals: CHASM’s SNVBOX [18], Mutation Assessor [25] and ANNOVAR [82]. Among them were conservation scores (such as ‘GERP, scores, ‘HMMPHC’, scores and others), amino acid substitution features (such as ‘PREDRSAE’, ‘PredBFactorS’, and others), exon features (such as ‘ExonSnpDensity’, ‘ExonConservation’, and others), features indicative of protein domain knowledge (such as “UniprotDOM_PostModEnz’, ‘UniprotREGIONS’, and others) and functional impact scores computed by algorithms such as VEST [23] and CHASM [18]. A tiny fraction (0.1%) of the UniProtKB annotations were not available from the SNVBOX database for our training data. We used the *k*-nearest neighbors-based imputation technique to substitute the missing features with those of the same gene’s nearest mutations. Our external validation datasets were free from any missing information.

### Density Estimation

A kernel density estimator (or KDE) takes an *n*-dimensional dataset as an input and outputs an estimate of the underlying *n*-dimensional probability distribution. A Gaussian KDE uses a mixture of *n*-dimensional Gaussian probability distributions to represent the density being estimated. It essentially tries to center one Gaussian component per data point, resulting in a non-parametric estimation of the density. One of the hyperparameters for a kernel density estimator is the bandwidth, which controls the kernel’s size at each data point, thereby affecting the “smoothness” of the resulting curve. We estimated the underlying probability distributions for the driver and passenger neighborhoods using a Gaussian kernel density estimator.

The schematic workflow of the entire process for a single run of the kernel density estimation experiment is shown in Figure 2(A-F). First, we randomly selected, with replacement, an equal number (*n*) of driver and passenger mutations from our training data for a single run of the kernel density estimation algorithm and particular window size (Figure 2A). Then, we tuned the bandwidth hyperparameter for each class of mutations using a 5-fold cross-validation approach and used the best parameters to derive the kernel density estimates (Figure 2B). Finally, we used the Jensen-Shannon (JS) distance metric to calculate the similarity between the two class-wise density estimates (Figure 2C). The JS distance between two probability distributions is based on the Kullback-Leibler (KL) divergence, but unlike KL divergence, it is bounded and symmetric. For two probability vectors, p and q, it is given by,

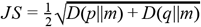

where 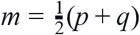, and *D* is the KL divergence. The significance of the estimated distances between the probability estimates was calculated using a randomized bootstrapping approach. Specifically, we randomly sampled with replacement twice the number (*2n*) of mutations from the same training set, irrespective of the labels. We then split the dataset in half, randomly assigning each half to driver and passenger mutations, respectively (Figure 2D). This was followed by a similar process of tuning the hyperparameters and deriving the class-wise density estimates (Figure 2E). Finally, we reported the JS distance between the density estimates (Figure 2F).

**Figure 2:**
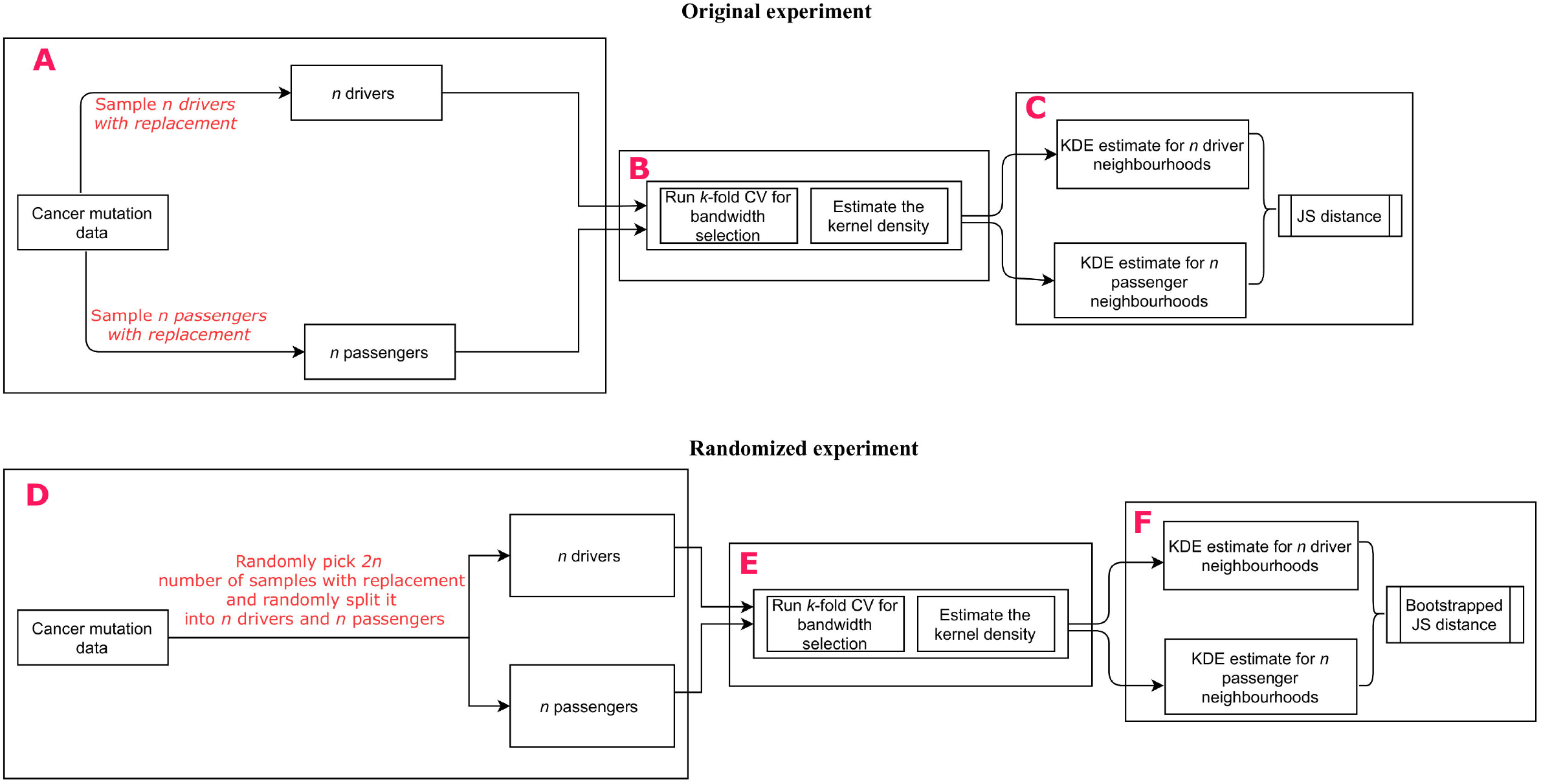
The workflow depicting one run of the kernel density estimation experiment is shown in this figure. All 5265 mutations from the Brown *et al*. study were used to derive the estimates. **(A)** First, an equal number of driver and passenger mutations were sampled with replacement. **(B)** The “bandwidth” hyperparameter was tuned using a 5-fold cross-validation approach, and the resulting tuned hyperparameter was used to estimate the densities. **(C)** The kernel density estimates for the driver and passenger neighborhoods were obtained separately, and the distance between them was calculated using the Jensen-Shannon (JS) distance. The JS distance is used to quantify how “distinguishable” two probability distributions are from each other. It is bounded between 0 and 1, where 0 represents the case where the two probability distributions are equal and vice versa. **(D)** The bootstrapping experiment to compute the significance of the density estimates calculated in (C) is shown in this figure. First, it involved random sampling of twice the driver or passenger mutations from (A) irrespective of the labels, followed by randomly splitting the data into driver and passenger labels. **(E)** Hyperparameter tuning and density estimation was performed similarly to (B). **(F)** The bootstrapped JS distance between the driver and passenger neighborhoods was derived. All six steps (A-F) of the density estimation experiments were repeated 30 times for all possible window sizes between 1 and 10 and seven different feature representations. The significance of the difference between the medians of the original and the bootstrapped JS distances was then reported.

We experimented with the following seven different neighborhood-based feature representations:

- One-hot encoding
- Count Vectorizer (*k-*mer sizes of 2,3 and 4)
- TF-IDF Vectorizer (*k-*mer sizes of 2,3 and 4)

The aforementioned KDE estimation experiments were repeated 30 times for all possible window sizes between 1 and 10 and all seven feature representations. Next, the best median JS distance estimate from the original experiments was reported for the given window size. The percentage of runs of the randomized experiments for which the estimated distance was greater than the original estimate was reported as the *p*-value.`The KernelDensity()` function from the scikit-learn *neighbors* module was used to derive the density estimates and `jensenshannon()` from the scipy *spatial.distance* submodule was used to calculate the distance metric.

### Classification Models

To build our binary classification models, we implemented three classifiers: the Random Forest classifier, the Extra Trees classifier (Extreme Random Forests), and the generative KDE classifier. The overall approach for the KDE-based classification was as follows (Figure 3A):

1. The dataset was split using the cross-validation strategy.
2. The training data was then split by label (driver/passenger).
3. For each class, we fit a generative model using the kernel density estimation method as described in the previous section. This gave us the likelihood that *P (x\passenger)* and *P (x\driver)* respectively for a particular data point *x*.
4. Next, the class prior, which is given by the number of examples of each class: *P (driver)* and *P (passenger)* was calculated.
5. Now, for a test data point *x*, the posterior probability was given by *P (driver\x)* ∞ *P (x\driver)P(driver)* and *P (passenger\x)* ∞ *P (x\passenger)P (passenger)*. The label that maximized the posterior probabilities was the one assigned to *x*.

**Figure 3:**
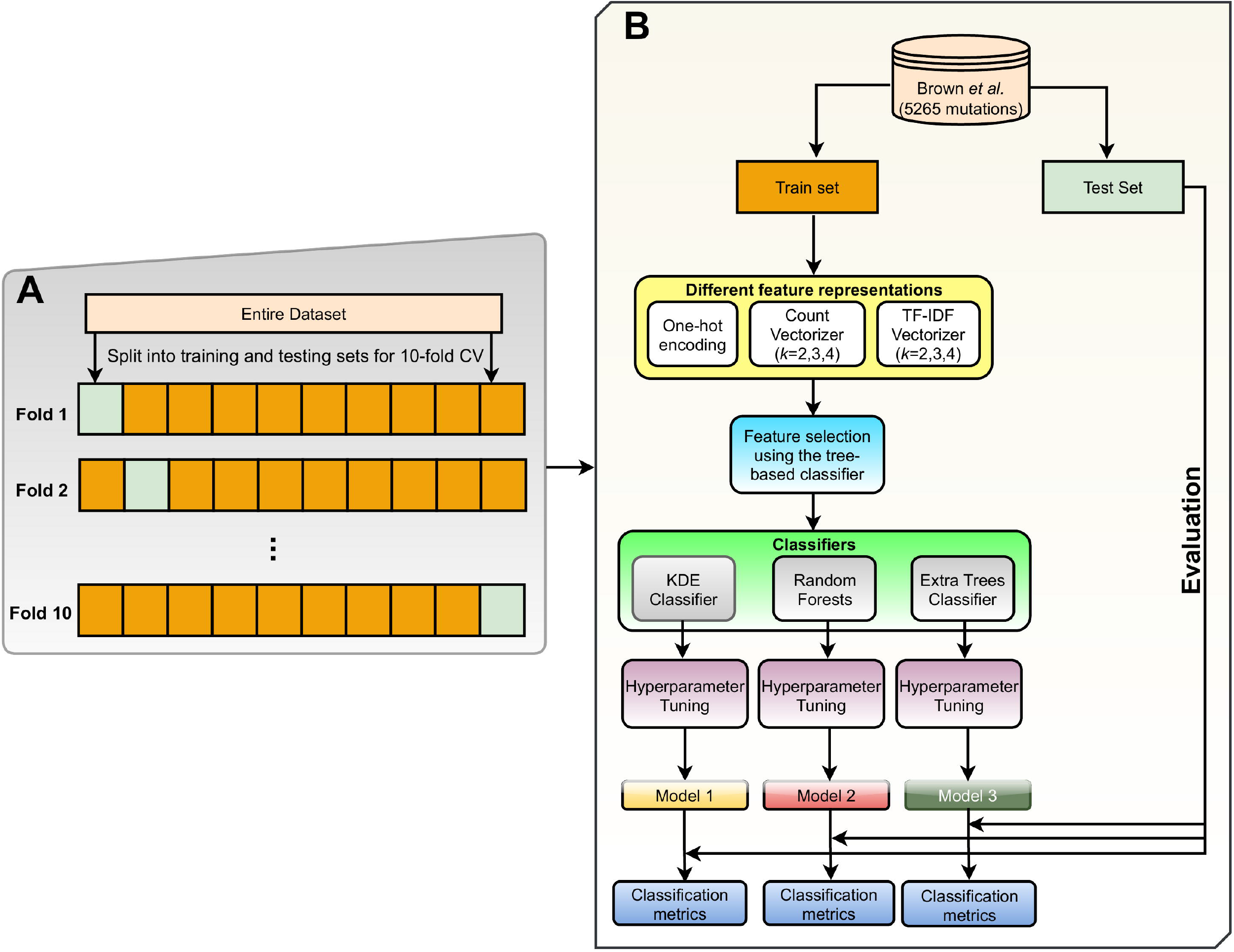
The workflow depicting one run of the 10-fold cross-validation experiments is shown in this figure. **(A)** In the first step, the entire dataset was split into ten equal parts. Nine of the ten subsets were combined into one training set, and one part was left as the test set. **(B)** Seven different feature representations [OHE, Count Vectorizer (*k*=2,3,4) and TF-IDF Vectorizer (k=2,3,4)] were considered for further analysis. After feature selection using a tree-based classifier, hyperparameter tuning was performed for three classifiers, and the corresponding models were derived. Finally, validation of each of the classifiers on the test set was performed, and the corresponding performance metrics were reported.

In contrast, both the tree-based classifiers are discriminative. They are composed of a large collection of decision trees where the final output is derived by combining every single tree’s predictions by a majority voting scheme. The main difference between the two tree-based classifiers lies in selecting splits or cut points to split the individual nodes. Random Forest chooses an optimal split for each feature under consideration, whereas Extra Trees chooses it randomly. All the classification models were written using the predefined functions available in the *scikit-learn (v. 0.22)* [83] module.

### Model Selection and Tuning

#### Repeated Cross-Validation Experiments

Owing to the relatively smaller sample size (5265 mutations) of the training set of mutations, we adopted a repeated 10-fold cross-validation approach to building our model. First, we split the dataset into ten equal subsets in a stratified fashion. Splitting the dataset in a stratified fashion maintains the same proportion of mutations in each class as observed in the original data. Nine of the ten subsets were combined into one training set (Figure 3A). In each training phase, we performed feature selection using the Extra Trees classifier, cross-validated grid search-based parameter tuning, training the classifiers using the best parameters, and obtaining the corresponding prediction scores on the hold-out test set (Figure 3B). For a given window size, we experimented with a total of seven feature representations (One-hot encoding, Count Vectorizer (*k-*mer size=2, 3 and 4), TF-IDF Vectorizer (*k*-mer size=2, 3 and 4), and three binary classifiers (Random Forests, Extra Trees, and Kernel Density Estimation). So overall, we had 21 distinct feature-classifier pairs.

We ran the 10-fold cross-validation experiments (Figure 3(A-B)) three times for each such pair, thereby obtaining 30 values for each classification metric: sensitivity, specificity, AUC, and MCC. The best overall median value, the 95% CI for each of the above metrics, and the corresponding feature-classifier pair were reported. To study the variation in classification performances with the addition of more nucleotides (or increase in window size), we repeated the Wilcoxon signed-rank test on the generated performance metrics for all 45 pairs of window sizes (*x,y*), where *x < y and (x,y)* ∈ [1,2,…, 10]. The `ci()`from the *gmodels* package [84] in R was used to calculate the 95% CIs for the various classification metrics.

#### Derivation of the Binary Classification Model to Distinguish between Driver and Passenger Mutations

To derive the final machine learning model, all overlapping mutations between the training set Brown *et al*., and the validation set Martelotto *et al*., were discarded, and the classifier was retrained on the reduced train set (4549 mutations: 544 drivers and 4005 passengers). The set of 989 mutations published by Martelotto *et al*. [43] formed our independent test set. Due to the inherent imbalance in the dataset, we implemented an undersampling technique known as Repeated Edited Nearest Neighbors [85] to downsize the majority class and consequently obtain a balanced dataset for subsequent training.

Predictions were obtained using two separate feature sets: 1) only neighborhood features based on the raw nucleotide sequences (or the neighborhood-only-model) and 2) neighborhood features plus the descriptive genomic features (or NBDriver). In addition to Random Forests, Extra trees, and the KDE classifier, we also experimented with a fourth classifier: a linear kernel SVM to obtain these predictions. Various combinations of these classifiers were implemented as ensemble models using the `VotingClassifier()` of the *ensemble* module in *scikit-learn*.

#### Feature Selection

We adopted an impurity-based feature selection technique for feature selection using the extra trees classifier to derive a ranked list of the top predictive features for our analysis. For the repeated cross-validation experiments, the features that were within the top 30 percentile of the most important features were selected and subsequently used to train our models. However, for deriving NBDriver, we built several classification models based on the top *n* (*n*=20, 30, 40, 50, 60) features and chose the one that gave the best overall classification performance.

The TF-IDF and Countvectorizer scores, used as features for our analysis, were implemented using the *feature_extraction* module in *scikit-learn*. In both cases, a new vocabulary dictionary of all the *k*-mers was first learnt from the training data using the `fit_transform()` routine and the corresponding term-document matrix was returned. Using this vocabulary, the scores of the *k*-mers from the test data were obtained using the `transform()` routine and were subsequently used in our analysis.

#### Hyperparameter Tuning and Classifier Threshold Selection

Hyperparameter tuning was done using a cross-validation based grid search technique over a parameter grid. The `GridSearchCV()` from the *model_selection* module in *scikit-learn* was used for this purpose. To further fine-tune the classifiers, we experimented with various classification thresholds from 0 to 1 with step sizes 0.001 and chose the one that gave the best AUROC. For an imbalanced classification problem, using the default threshold of 0.5 is not a viable option and often results in the incorrect prediction of the minority class examples.

#### Performance Metrics

For the repeated cross-validation experiments, we assessed our classifiers’ performance using four commonly used performance metrics: Sensitivity, Specificity, Mathews correlation coefficient (MCC), and Area under the ROC curve (AUROC). Mathews correlation coefficient is a balanced metric and is very useful in imbalance classification problems. It is bounded between −1 and 1, with −1 representing perfect misclassification, 0 representing average classification, and +1 representing ideal classification. It is given by the following expression:

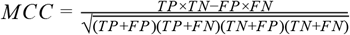

where TP stands for True Positives, TN, True Negatives, FP, False Positives and FN, False Negatives. MCC is a more robust alternative to Accuracy and F1-score that can sometimes show overoptimistic classification performance for imbalanced data and was therefore not used for the analysis.

After deriving the binary classifier, we used additional classification performance metrics outlined by Martelotto *et al*. to compare our algorithm’s performance with other state-of-the-art mutation effect prediction tools. They were Positive Predictive Value (PPV), Negative Predictive Value (NPV), and a composite score, defined as the sum of Sensitivity, Specificity, PPV, and NPV.

#### Comparison with Other Pan-Cancer Mutation Effect Predictors

Similar to the benchmarking study conducted by Martelotto *et al*., we compared the generated binary classifiers with nine pan-cancer mutation effect prediction tools: Mutation Taster [86], FATHMM (cancer) [19], Condel [26], FATHMM (missense) [19], PROVEAN (v1.1.3) [16], SIFT (Ensemble 66) [87], Polyphen2 [17], Mutation Assessor [25] and VEST [23] using the set of 989 literature-curated mutations. For each of these predictors, we used the prediction labels based on predefined score cutoffs published as part of the Martelotto *et al*. [43] study. Two new prediction algorithms (CHASMplus (pan-cancer) [24] and CanDrA+ (Cancer-in general) [27]) were also added to the list, and the score cutoffs were decided in the following manner.

For CHASMplus, we tested all possible thresholds between 0 and 1 with step sizes of 0.01 and chose the corresponding threshold with the highest composite score due to the absence of a default threshold. All mutations with predicted scores greater than this optimal threshold were labeled as drivers and vice versa. For CanDrA+, we used the default prediction categories [27]. Predictions for CHASMplus and CanDrA+ were obtained from the OpenCRAVAT web server [88] and executable packages published by Mao *et al*. [27]. Different mutation effect predictors were combined using the majority voting rule to obtain better predictive power, and ensemble models were created.

While comparing two algorithms, to derive the significance of the difference between any two classification metrics, we adopted the same strategy as Martelotto *et al*. Briefly, we derived the 95% CI for each of these classification metrics by repeated sampling with replacement with 1000 iterations. If the generated CI’s touched or there was no overlap, the difference was considered significant (*p* <0.05) based on the results of the analysis done by Ng *et al*. [89].

## Results

First, we report a pan-cancer machine learning tool, NBDriver (**N**eighborhood **D**river), which utilizes neighborhood sequences as features to discriminate missense mutations as either drivers or passengers. Our key results are three-fold. First, we use generative models to derive the distances between the underlying probability estimates of the two classes of mutations. Then, we build robust classification models using repeated cross-validation experiments to derive the median values of the metrics designed to estimate the classification performances. Finally, we demonstrate our models’ ability to predict unseen coding mutations from independent test datasets derived from large mutational databases.

### Neighborhood Sequences of Driver and Passenger Mutations Show Markedly Different Distributions

We estimated the driver and passenger neighborhood sequences’ underlying probability distributions using kernel density estimation. We computed the Jensen–Shannon (JS) distance metric to understand how “distinguishable” they are from one another. The JS metric is bounded between 0 (maximally similar) and 1 (maximally dissimilar). Table 3 shows the results of the KDE estimation experiments for various window sizes. We observed that, for the Brown *et al*. dataset [37]‘ the maximum significant (*p* < 0.05) median JS distance between passenger and driver neighborhood distributions, calculated across 30 runs of bootstrapping experiments, was 0.275 (for a window size of 2), and the minimum was 0.211 (for window sizes 7-10). Figure 4 shows the variation in the JS distances between the original and the randomized KDE experiments for window sizes between 1 and 10. As evident from Figure 4, except for window size 1, all other window sizes had a significant JS distance value (*p* < 0.05).

**Table 3:**
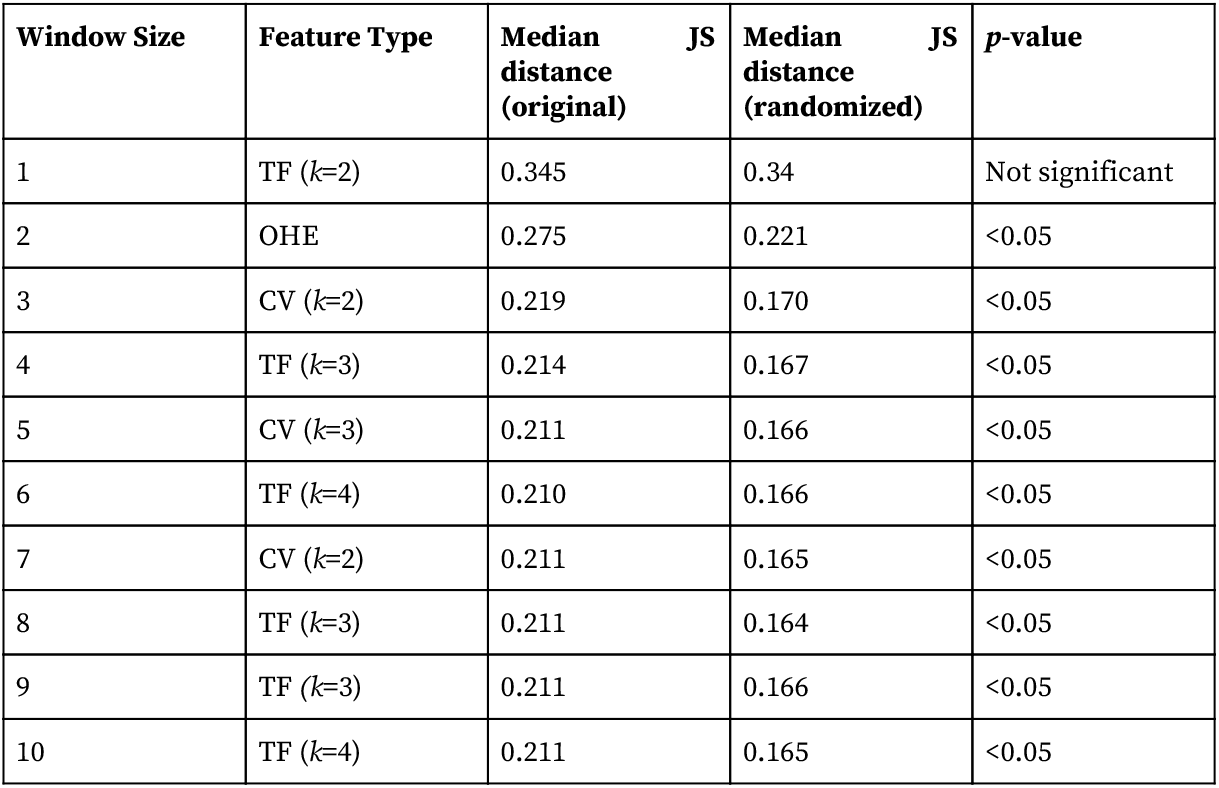
Median JS distances for both the original and randomized experiments for different window sizes

**Figure 4:**
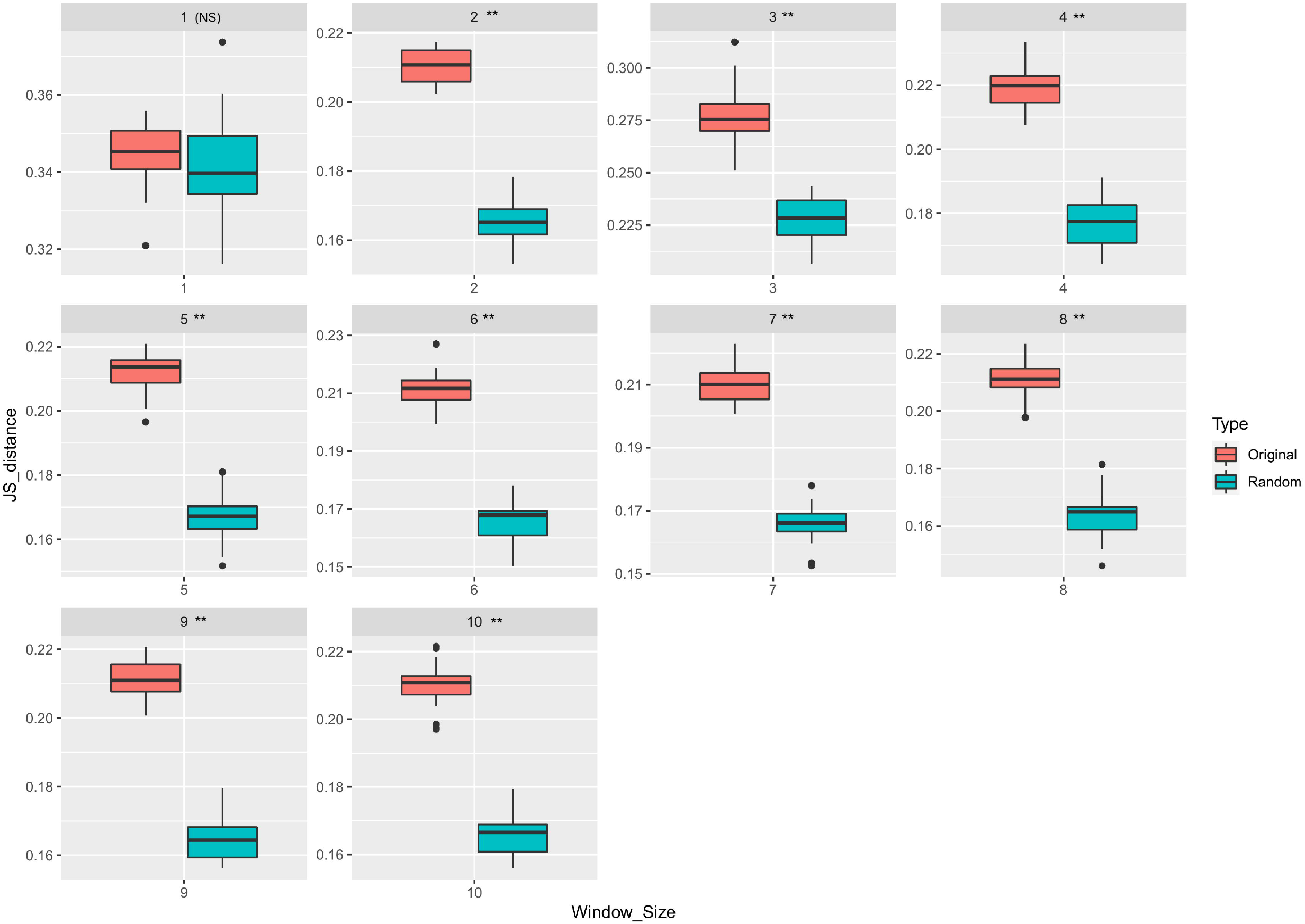
Variation in JS distances between the estimated densities for every window size between 1 and 10 is shown in this figure. All 5265 mutations from the original study were used here. Two types of boxplots, one for the original and another for the randomized experiments have been shown here along with the p-values, which approximates the probability that the original median distance can be obtained by chance. Except window 1, all other window sizes had a significant (** *p* < 0.05) difference between the original and the randomized JS distances.

Out of the seven different feature representations, we reported the ones that gave the maximum median JS distance. From Table 3, we observed that a TF-IDF vectorizer with *k*-mer sizes 2,3 and 4 was the preferred form of feature representation for six window sizes (1, 4, 6, 8, 9 and 10), whereas a count vectorizer with *k*-mer sizes 2 and 3 was chosen for three window sizes (3, 5, and 7). However, the only exception was for a window size of 2, where the one-hot encoding-based feature representation technique gave the maximum median JS distance. These results indicated the TF-IDF based feature representation was the most efficient at delineating the differences in the distributions between the driver and passenger neighborhoods.

### Repeated Cross-Validation Using Neighborhood Features Generates Robust Classification Models

The repeated cross-validation experiments using only the neighborhood sequences as features are shown in the Supplementary Table 3A. From these results, we observed that the best median sensitivity of 0.938 (95%CI 0.919-0.940) was obtained using features derived from a count vectorizer and subsequent training using a random forest classifier for window sizes 1, 5, 6 and 9. However, the best median specificity of 0.807 (95%CI 0.791-0.811), AUC of 0.832 (95% CI 0.826-0.841), and MCC of 0.584 (95% CI 0.564-0.594) were obtained using a TF-IDF based feature representation trained using a KDE classifier for a window size of 10. The variation in the classification performances for different window sizes obtained during the repeated cross-validation experiments using the initial training set of 5265 mutations is shown in Figure 5. This figure shows that except for window sizes 1 and 2, a TF-IDF vectorizer gave the maximum median AUC, Specificity, and MCC. However, for all window sizes, the maximum median sensitivities were obtained using the count vectorizer based feature representation technique. Classification metrics such as AUC and MCC are used to measure the quality of binary classifications. Similar to our observations made from the KDE estimation experiments (Table 3), the TF-IDF vectorizer performed consistently well both in terms of the overall AUC and MCC, indicating that this particular feature representation technique was the most efficient separating the two classes of mutations.

**Figure 5:**
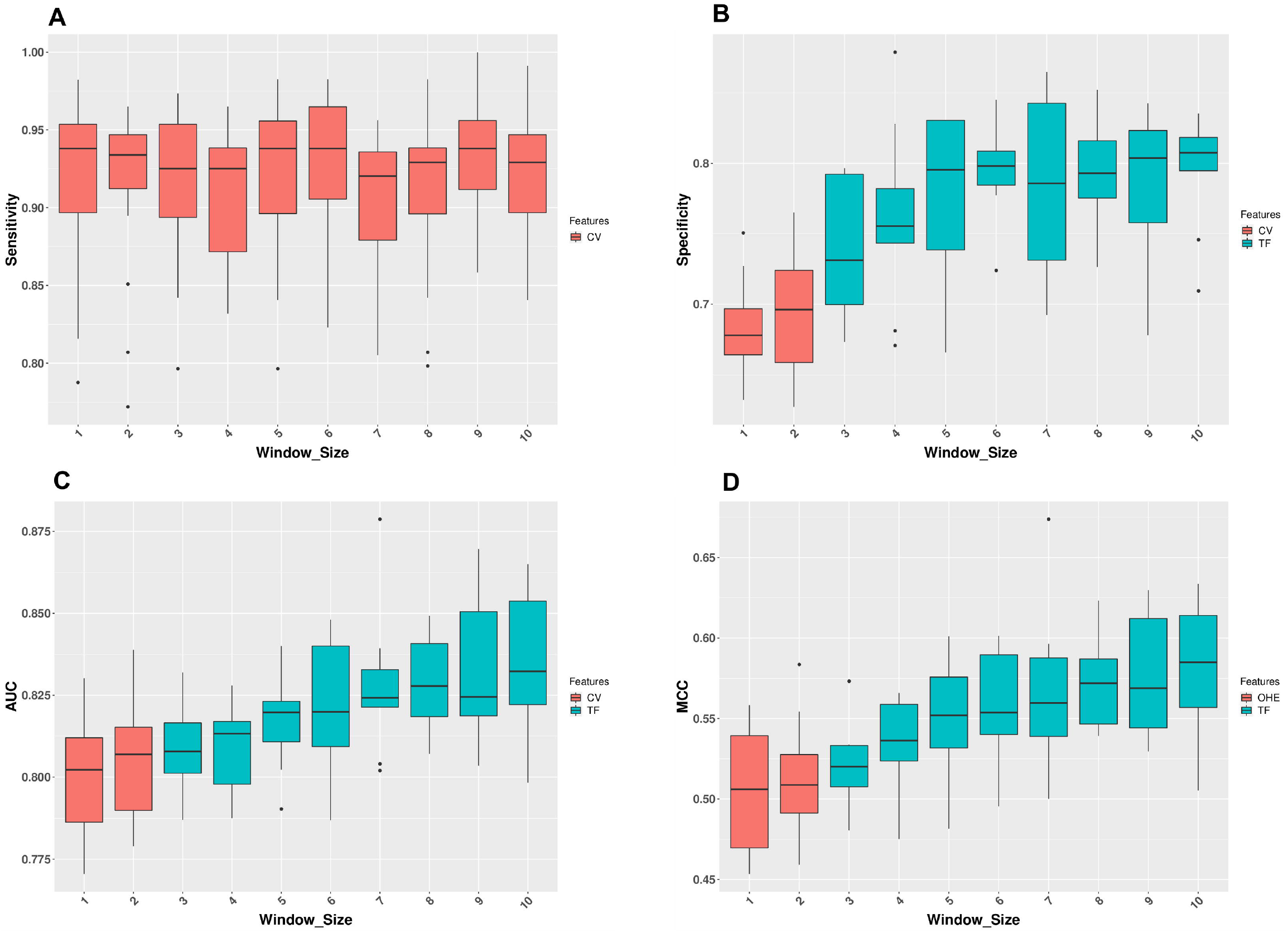
The variation in the classification performances with different window sizes obtained during the repeated cross-validation experiments using the initial training set of 5265 mutations is shown in this figure. For each window size, feature representations among CV (CountVectorizer), TF (TF-IDF Vectorizer) and OHE (One-hot encoding) that gave the best performances in terms of **(A)** Sensitivity **(B)** Specificity **(C)** AUC and **(D)** MCC is displayed.

The variation in the classification performances with the increase in the window size is shown in Supplementary Table 3b. From this table, we observed that out of the 45 unique pairs of window sizes (Methods: Repeated cross-validation experiments), 27 had a significant (*p* < 0.05; Wilcoxon signed-rank test) increase in specificity and AUC while 31 had a significant (*p* < 0.05; Wilcoxon signed-rank test) increase in MCC with the addition of more nucleotides. However, for sensitivity, a significant increase was observed only when the window size was increased from 4 to 9 and 7 to 9, respectively. These results indicated that adding more nucleotides to a particular window does not always guarantee an increase in the classifier’s performance in distinguishing between driver and passenger mutations.

### Classification Models Give Performances Comparable with Other State-of-the-Art Mutation Effect Predictors

Using only the neighborhood nucleotide sequences as features, the best results (Table 4A) on the independent test set [38]‘ was obtained using an Extra Trees classifier. This neighborhood-only model was trained on features extracted using the Count Vectorizer technique on a window size of 10.

**Table 4a:** Comparison of the generated binary classifiers with other mutation effect prediction algorithms using the benchmarking dataset by Martelotto *et al*.

| Algorithm | Accuracy | Sensitivity | Specificity | PPV | NPV | CS | MCC |
| --- | --- | --- | --- | --- | --- | --- | --- |
| Mutation Taster | 0.8857 | 0.9081 | 0.75 | 0.9566 | 0.5738 | 3.1885 | 0.590 |
| FATHMM (Cancer) | 0.91 | 0.9788 | 0.4929 | 0.9213 | 0.7931 | 3.1861 | 0.580 |
| CHASMplus<br>(Pancancer) | 0.85 | 0.852 | 0.85 | 0.972 | 0.486 | 3.16 | 0.570 |
| **NBDriver** | 0.891 | 0.931 | 0.643 | 0.941 | 0.608 | 3.123 | 0.561 |
| **Neighborhood-only**<br>**model** | 0.85 | 0.629 | 0.907 | 0.9744 | 0.285 | 2.7954 | 0.370 |
| Condel | 0.8584 | 0.9258 | 0.45 | 0.9108 | 0.5 | 2.7866 | 0.392 |
| FATHMM (missense) | 0.8251 | 0.8775 | 0.5071 | 0.9152 | 0.4057 | 2.7055 | 0.351 |
| PROVEAN | 0.7371 | 0.7444 | 0.6929 | 0.9363 | 0.3089 | 2.6825 | 0.327 |
| SIFT | 0.8099 | 0.861 | 0.5 | 0.9126 | 0.3723 | 2.6459 | 0.32 |
| Polyphen-2 | 0.7978 | 0.8422 | 0.5286 | 0.9155 | 0.3558 | 2.6421 | 0.317 |
| Mutation Assessor | 0.747 | 0.7665 | 0.6286 | 0.9259 | 0.3077 | 2.6287 | 0.3 |
| VEST | 0.7503 | 0.8269 | 0.2857 | 0.8753 | 0.2139 | 2.2018 | 0.1 |
| CanDrAplus<br>(Cancer-in-general) | 0.592 | 0.857 | 0 | 0.99 | 0 | 1.847 | -0.03 |

We trained NBDriver by combining the neighborhood features and the descriptive genomic features. Out of the various classifiers implemented, an ensemble model consisting of a linear kernel SVM and a KDE classifier gave the best results (Table 4A). Compared to the neighborhood-only model, there was a significant increase (*p* < 0.05) in accuracy (=0.891), sensitivity (=0.93), NPV (=0.608), Composite Score (=3.123), and MCC (=0.561). However, this was accompanied by a significant (*p* <0.05) drop in specificity (=0.643). There was no significant change in PPV, though.

A ranked list of the 50 features used to train NBDriver is shown in Supplementary Table 4. Out of those 50 features, 26 were neighborhood-based features or the TF-IDF scores of the overlapping 4-mers extracted from a window size of 10. The plot displaying the variation in the AUROC with various classification thresholds is shown in Figure 6. The best results were obtained using a threshold of 0.119. Consequently, all mutations with the prediction scores above this threshold were classified as drivers and vice versa.

**Figure 6:**
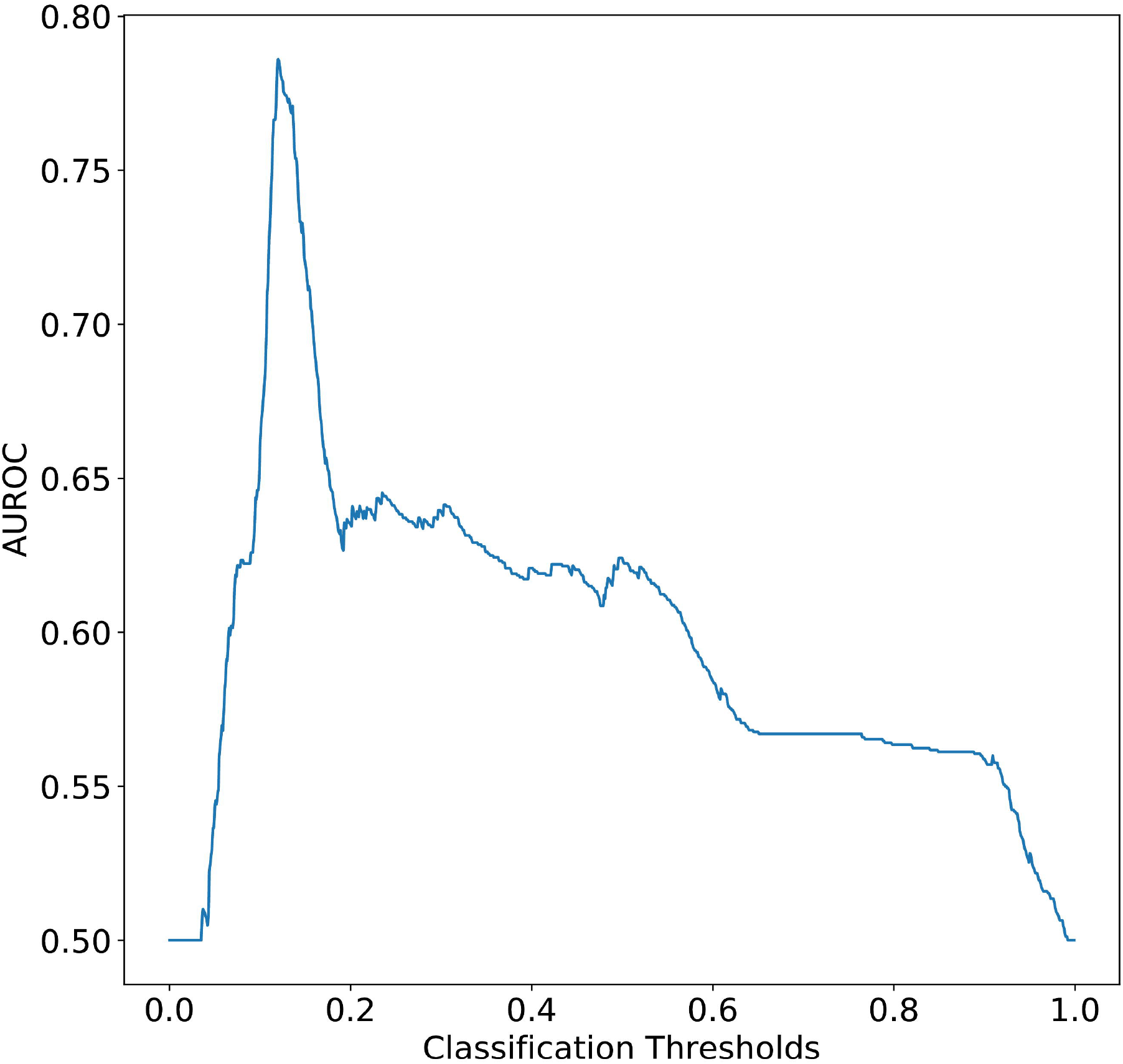
Plot showing the variation in AUROC with the different classification thresholds obtained while deriving NBDriver is shown here. NBDriver was trained on a reduced training set of 4549 mutations after removing all overlapping mutations from the original study and Martelotto *et al*. For an imbalanced classification problem, using the default threshold of 0.5 is often not advisable. In our case, the best AUROC was obtained using a threshold of 0.119. Consequently, all mutations with prediction scores greater than this threshold were classified as drivers and vice versa.

Overall, on this benchmarking dataset, NBDriver ranked fourth in terms of the composite score, fifth in terms of specificity, and second in NPV, PPV, Sensitivity, and Accuracy. By contrast, although the neighborhood-only-model was the top-ranking tool in terms of Specificity and PPV, it did not perform well in terms of the other metrics. Owing to NBDriver’s superior performance, all subsequent external validations were performed using this model.

### Voting Ensemble of Prediction Algorithms Gives Better Classification Performances

We also assessed the effect of combining multiple top-ranked single predictors into an ensemble model. We evaluated NBDriver’s contribution to the overall ensemble by obtaining predictions without the tool. The top-performing ensemble consisting of NBDriver, CHASMplus, FATHMM (cancer), Mutation Taster, and Condel resulted in a composite score of 3.504, accuracy of 0.945, and an NPV of 0.88, significantly higher (*p* < 0.05) than every single predictor evaluated in the study (Table 4b; Supplementary Table 5). The composite score and accuracy obtained using this ensemble were also the highest among all the different combinations of single-predictors tested in this study (Supplementary Table 5). Removing NBDriver from the ensemble resulted in a significant decrease (*p* < 0.05) in the composite score, NPV, MCC, Accuracy, and Sensitivity. However, it was accompanied by a significant increase in specificity and no significant PPV change for the smaller ensemble (Table 4b).

**Table 4b:** Evaluating the contribution of NBDriver to the top performing ensemble

| Algorithm | Accuracy | Sensitivity | Specificity | PPV | NPV | CS | MCC |
| --- | --- | --- | --- | --- | --- | --- | --- |
| **NBDriver** +<br>CHASMplus+<br>FATHMM (cancer) +<br>Mutation Taster +<br>Condel | 0.945 | 0.985 | 0.689 | 0.95 | 0.88 | 3.504 | 0.746 |
| CHASMplus+<br>FATHMM (cancer) +<br>Mutation Taster +<br>Condel | 0.921 | 0.942 | 0.771 | 0.96 | 0.71 | 3.384 | 0.691 |
| A smaller ensemble that gave no significant change in Composite score and MCC compared to the previous ensemble (First row of Table 4b) |  |  |  |  |  |  |  |
| **NBDriver** + Mutation<br>Taster + Condel | 0.942 | 0.99 | 0.65 | 0.945 | 0.919 | 3.504 | 0.745 |

Another ensemble model consisting of NBDriver, Mutation Taster, and Condel gave similar results (Composite score=3.504) as the previous one (Table 4b; Supplementary Table 5). Compared to the previous ensemble (Table 4b), there was no significant difference in MCC, Composite Score, PPV, Sensitivity, and Accuracy. However, there was a significant increase in the NPV and a significant decrease in the specificity.

A complete set of all the different combinations of the single predictors evaluated in this study is present in Supplementary Table 5. From this table, we observed that the maximum sensitivity (=0.9941) and NPV (=0.9375) were obtained by the ensemble (Mutation Taster, FATHMM (cancer), and CONDEL), which did not include NBDriver. However, the maximum specificity (=0.8357) and PPV (=0.9711) were obtained using the ensemble (NBDriver, CHASMplus, Mutation Taster, and CONDEL).

### Driver and Passenger Mutations, Features Used to Train NBDriver are Significantly Different

Our feature selection results illustrate the differences in the underlying biological processes governing driver and passenger mutations similar to Mao *et al*. [27]. Using the training data used to build NBDriver, we found that driver mutations tend to occur on amino acid residues that have stiff backbones and have less solvent accessibility as denoted by the significantly lower (Wilcoxon test; *p* < 5.4 × 10^-10^) ‘PREDRSAE’ probability measure (Figure 7A) and the significantly higher (Wilcoxon test; *p* < 2.1 × 10^-9^) ‘PredBFactorS’ probability measure (Figure 7B) respectively. We also observed that a mutation is more likely to be a driver if it occurs in genomic regions that were evolutionarily conserved. The mean GERP score for driver mutations was significantly higher (Wilcoxon test; *p* < 2.2 × 10^-16^) than that of passengers (Figure 7C). Similarly, driver mutations were more common in genomic sites that had a significantly higher (Wilcoxon test; *p* < 3.3 × 10^-16^) Positional Hidden Markov Model (HMM) conservation score (or HMMPHC) as compared to passengers (Figure 7D). Among the other features, we observed similar class-wise distributional differences among features indicative of protein domain knowledge. ‘UniprotDOM_PostModEnz’ denotes the presence or absence of a mutation in a site within an enzymatic domain responsible for post-translational modification (or PTM). PTM-related mutations are often accountable for changes in protein functions and alterations of regulatory pathways, eventually leading to carcinogenesis. ‘UniprotREGIONS’ is another binary feature that tells us whether a mutation occurred in an experimentally defined region of interest in the protein sequence, such as those associated with protein-protein interactions and regulation of biological processes. Our analysis pointed out that a considerable portion (31%) of driver mutations clustered around PTM sites, contrasted by around 0.4% of passengers (Figure 7E). Similarly, about 37% of driver mutations were located in protein domains that were experimentally defined as regions of interest compared to around 11% of passengers (Figure 7F).

**Figure 7:**
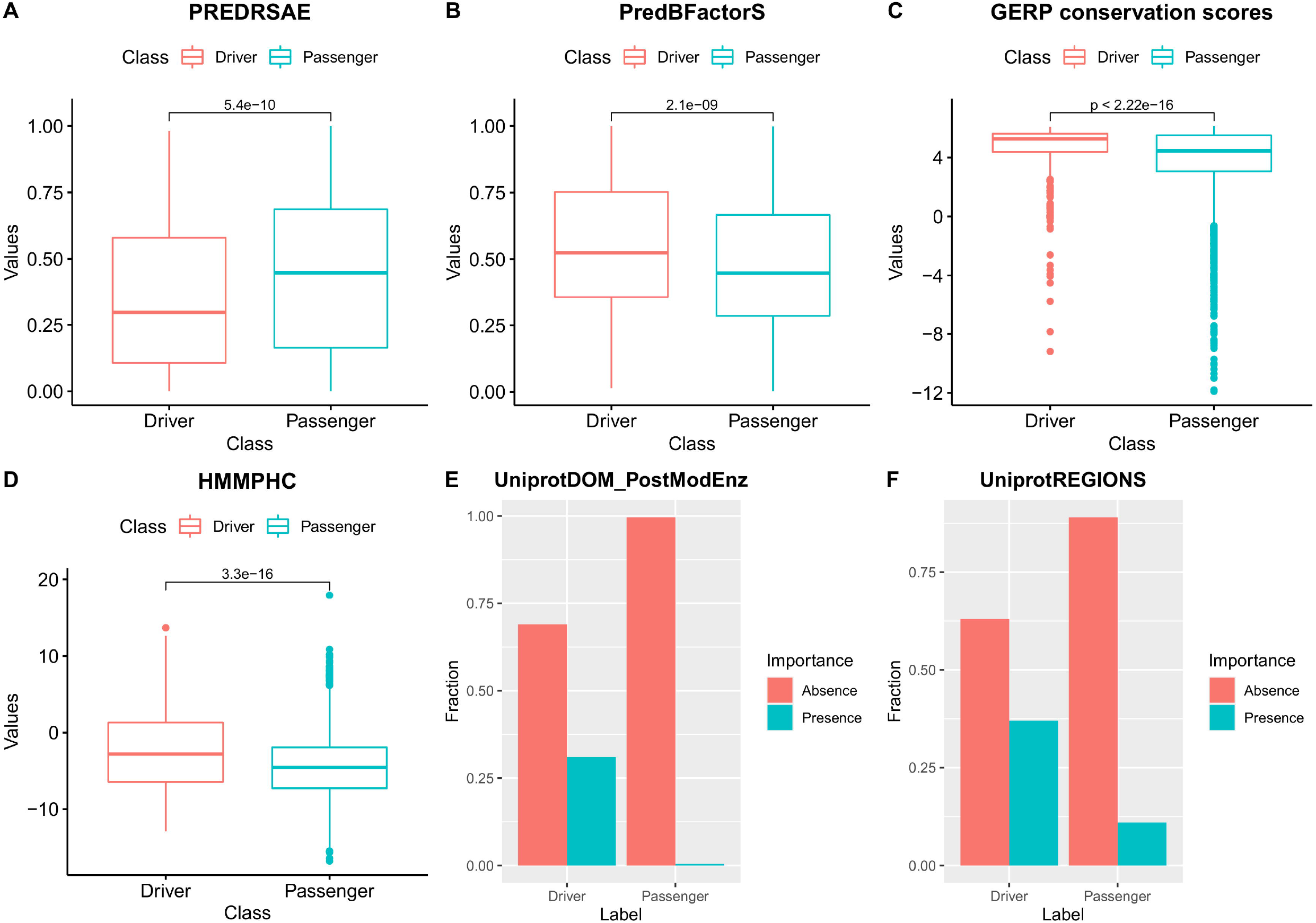
Differences in the distribution of features between driver and passenger mutations observed from the training data used to derive NBDriver. **(A) PREDRSAE** (Predicted Residue Solvent Accessibility - Exposed) gives the probability of the wild type residue being exposed. From the plot it is clear that probability of driver mutations occurring in residues that are exposed is significantly less (Wilcoxon test; *P*=5.4E-10) than that of passengers. **(B) PredBFactorS** (High Predicted Bfactor) gives the probability that the wild type residue backbone is stiff. From the plot it is clear that the probability of driver mutations occurring in residues with stiff backbones is significantly higher (Wilcoxon test; *P*=2.1E-09) than that of passengers. **(C) GERP conservation scores** give the evolutionary conservativeness scores for specific sites where mutations have occurred. From the plot it is clear that driver mutations occur in sites with GERP scores that are significantly higher (Wilcoxon test; *R*<2.2E-16) than passenger mutations. **(D) HMMPHC** (Positional Hidden Markov Model (HMM) conservation score) is a measure which is calculated on the basis of the degree of conservation of the residue, the mutation and the most probable amino acid. From the plot it is clear that driver mutations tend to occur in residues with HMMPHC scores significantly higher (Wilcoxon test; *P*=3.3E-16) than passenger mutations. **(E) UniprotDOM_PostModEnz** is a feature based on protein domain knowledge which tells us whether a site in an enzymatic domain is responsible for any kind of post translational modification (or PTM). ‘Presence’ indicates that the mutation occurs in a site responsible for PTM and vice versa. From the plot it is clear that more driver mutations occur in PTM-associated sites as compared to passengers. **(F) UniprotREGIONS** is a binary variable which tells us whether a mutation occurs in a region of interest in the protein sequence. ‘Presence’ indicates that the mutation occurs in a region of interest and vice versa. From the plot it is clear that more driver mutations cluster in regions of interest in the protein sequence as compared to passengers thereby making them mechanistically influential for the progression of the disease.

In our approach, the TF-IDF algorithm was used to weigh a *k*-mer and assign importance to it in the given set of neighborhood sequences. Also, a higher TF-IDF score is indicative of the greater relevance/importance of that *k*-mer. Our feature selection results indicated that for the 26 neighborhood sequence-based features, the mean TF-IDF scores for drivers were significantly higher (Wilcoxon test;*p* <0.05) than that of passengers (Figure 8). This result suggested that NBDriver’s top neighborhood features are more specific to the driver neighborhoods than the passengers.

**Figure 8:**
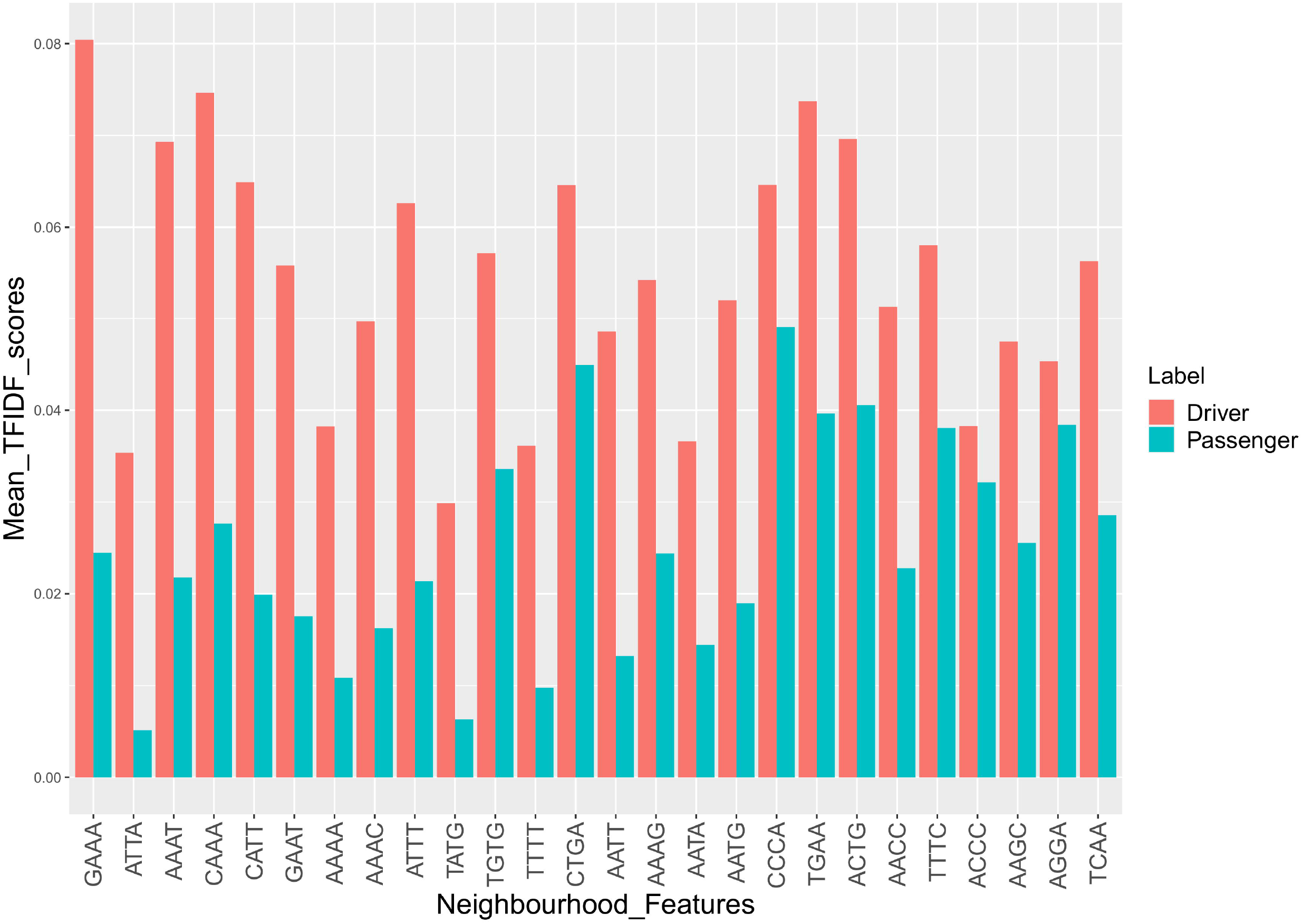
Plot showing the class-wise variation in the mean TF-IDF scores for the 26 neighborhood-sequence features used to train NBDriver. The x-axis represents the 4-mers used in the analysis, and the y-axis represents the mean TF-IDF scores. From the plot, it is evident that the mean TF-IDF scores are consistently higher for drivers as compared to passengers. Since a higher TF-IDF score indicates the relevance or importance of a particular *k*-mer, we can conclude that the 4-mers used to derive NBDriver are more specific to the driver neighborhoods than passengers.

### Evaluation Using Previously Unseen Coding Mutation Data

To evaluate NBDriver’s capability at identifying previously unseen driver mutations, we evaluated it using missense mutation data compiled from the following four databases.

#### Cancer Mutation Census

Based on the various evidence criteria set forth by the Cancer Mutation Census database, a particular mutation can be classified into tier 1, 2, or 3, with tier 1 mutations having the highest level of evidence of being a driver and so on. From the list of missense mutations in the CMC not present in our training data, NBDriver could accurately predict all 19 tier 1, 25 out of 28 tier 2, and 179 out of 230 tier 3 mutations, achieving an overall accuracy of 81%. On the other hand, the ensemble model consisting of NBDriver, Condel and MutationTaster could accurately predict all 19 tier 1, 27 out of 28 tier 2, and 214 out of 230 tier 3 mutations achieving an overall accuracy of 94%. Upon further investigation, we found that NBDriver was highly successful in identifying hotspot mutations present in the CMC. Recurrent alterations at the same genomic site in cancer genes such as MET, MPL, FLT3 and KIT have been implicated in many different cancer types [39]–[43] (Supplementary Table 7a).

#### Cancer Genome Interpreter Database

Using pathogenic mutations compiled from various sources, we found that NBDriver could accurately identify 1274 out of 1628 non-overlapping missense driver mutations, achieving an overall accuracy of 78%. The model correctly identified all three mutations from the Cancer Biomarkers Database, 39 out of 47 mutations from the DoCM database, 23 out of 31 mutations from the Martelotto *et al*. study [38], and 1209 out of 1547 mutations from the OncoKB database. On the other hand, the ensemble model comprising NBDriver, Condel and MutationTaster could accurately predict 1519 out of 1628 mutations achieving an overall accuracy of 93%.

#### Recurrent Driver Mutations

Out of the top 33 hotspot mutations identified in the study conducted by Rheinbay *et al*. [44] as recurrently mutated, NBDriver correctly identified 27 as drivers. However, Mutation Taster displayed superior performance by identifying all 33 mutations correctly. Except for KRAS, NBDriver correctly identified all mutations from the other four genes (NRAS, TP53, PIK3CA, and IDH1) as cancer drivers. Hotspot mutations in these four genes reported by Rheinbay *et al*. [44], correctly identified as drivers by NBDriver have been implicated in many different cancers [45]–[48] (Supplementary Table 7a).

#### Rare Driver Mutations Found in Glioblastoma and Ovarian Cancer

Using the list of rare drivers reported by the developers of the driver prediction tool CanDrA [27], we evaluated NBDriver’s ability to identify less frequent alterations in the cancer genome. Overall, NBDriver alone could identify 29 out of 34 (85%) glioblastoma mutations and 20 out of 38 (53%) ovarian cancer mutations. All these mutations belonged to eight known OVC-related genes (ARID1A, CDK12, ERBB2, MLH1, MSH2, MSH6, PIK3R1, PMS2) and seven known GBM-related genes (ATM, EGFR, MDM2, NF1, PDGFRA, PIK3CA, ROS1). All eight OVC-related genes correctly identified as drivers by NBDriver have been implicated in ovarian cancer through observations made from multiple studies [49]–[53] (Supplementary Table 7b). The ensemble model made up of NBDriver, Condel and Mutation Taster performed better than the single predictor by identifying 32 out of 34 (94%) glioblastoma mutations and 24 out of 38 (63%) ovarian cancer mutations.

#### Stratification Of the Predicted Driver Genes Based on Literature Evidence

We combined the list of genes with at least one true positive missense driver mutation prediction from NBDriver into a catalog of 138 putative driver genes. We then compared our gene set against those already published in six landmark pan-cancer studies for driver gene identification. Bailey *et al*. [54] identified 299 driver genes from 9423 tumor exomes by combining the predictions from 26 different computational tools. Martincorena *et al*. [55] used the normalized ratio of non-synonymous to synonymous mutations (dN/dS model) to identify driver genes from 7664 tumors and reported a total of 180 putatively positively-selected driver genes and 369 known cancer genes from three main sources:

1. 174 cancer genes from the version 73 of the COSMIC database [6].
2. 214 significantly mutated genes across 4742 tumors identified by Lawrence *et al*. [56] using the MutSigCV tool.
3. 204 genes identified through a literature search.

Two marker papers from TCGA [57], [58] identified 132 significantly mutated genes using the MutSigCV tool. Tamborero *et al*. [35] identified a list of 291 high-confidence drivers from 3205 tumor samples using a rule-based approach. Deitlein *et al*. [59] modelled the nucleotide context around driver mutations and identified 460 driver genes based on nucleotide context. Apart from the aforementioned studies, overlap between our list of genes and two well-established cancer gene repositories: the Cancer Gene Census [6], [60] and the Intogen database [61] was also reported. We identified 124 (=89%) of our predicted driver genes as canonical cancer genes present in the Cancer Gene Census. Among the remaining genes, six were catalogued as drivers in at least two of the pan-cancer studies or mutation databases as mentioned above (Supplementary Table 6). A total of eight genes (CTLA4, IGF1R, PIK3CD, TGFBR1, RAD54L, SHOC2, CDKN2B and XRCC2) were not identifiable from any of the landmark studies or databases and required further validation.

## Discussion

Our investigation aimed to compare the raw neighborhood sequences of driver and passenger mutations and exploit any observed distributional differences to build robust classification models. We showed that except for one window size (n=1), a significant difference in the distributions between the neighborhoods of driver and passenger mutations was consistently present in our cohort. Using TF-IDF and Count Vectorizer scores derived from the overlapping *k*-mers, we trained a KDE-based generative classifier and two other tree-based classifiers. One crucial distinction between NBDriver and other methods is the inclusion of overlapping *k*-mers extracted from the neighborhood of mutations as features for further analysis. NBDriver was trained using a small set (=50) of highly discriminative features, 52% of which were neighborhood scores. Using this model, we could accurately predict 89% of all the literature-curated mutations outlined in the Martelotto *et al*. study [38], 81% of the high confidence list of mutations recently published by the Cancer Mutation Census, 78% of all the actionable alterations reported in the Cancer Genome Interpreter, 82% of all the hotspot mutations reported from a pan-cancer genome analysis, 85% and 53% of rare driver mutations found in glioblastoma and ovarian cancer respectively. Ensemble models obtained by combining the predictions from other state-of-the-art mutation effect predictors with NBDriver performed significantly better than the individual predictors in all five validation datasets. These results underscore the importance of including neighborhood features to build mutation effect prediction algorithms.

Although our method’s focus was to identify missense driver mutations from sequenced cancer genomes, the majority of the genes (130 out of 138) containing at least one predicted mutation belonged to the Cancer Gene Census or other large-scale driver gene discovery studies. The protein products of the eight remaining genes not flagged as drivers by any of the databases/studies had known functional roles in maintaining the cancer genome’s stability and promoting tumor development. The CTLA4 gene modulates immune response by serving as checkpoints for T-cell activation, essentially decreasing the T cells’ ability to attack cancer cells. Immune checkpoint inhibitors, which are designed to “block” these checkpoints have drastically changed the treatment outcomes for several cancers [62]. Transcriptomic profiling of blood samples drawn from cervical cancer patients identified IGF1R as a biomarker for increased risk of treatment failure [63]. Overexpression of the PIK3CD gene has been associated with cell proliferation in colon cancer and is responsible for poor prognosis among patients [64]. Multiple studies have indicated an association with polymorphisms observed in TGFBR1 and cancer susceptibility [65], [66]. Similarly, polymorphisms detected in the RAD54L is a genetic marker associated with the development of meningeal tumours [67]. SHOC2 has been reported to be a regulator of the Ras signalling pathway and is associated with poor prognosis among breast cancer patients [68]. Similarly, the inactivation of the CDKN2B gene is responsible for the progression of pancreatic cancer [69]. With the help of massively parallel sequencing studies, rare mutations in the XRCC2 gene have been linked to increased breast cancer susceptibility among patients [70].

Our study does have some limitations. First, we used a representative dataset of driver and passenger mutations whose labels were not *in silico* predictions from other mutation effect prediction algorithms but derived from experimentally validated functional and transforming impacts from various sources. This resulted in a relatively small sample size for supervised classification. However, this approach also minimized the chances of inadvertently introducing false-positive mutations into the training set used to derive the driver and passenger neighborhoods’ class-wise density estimates or the machine learning models. Evidence [71] suggests that a sizeable proportion of mutations present in large mutational databases are mostly false positives, reflecting sequencing errors due to DNA damage. Moreover, NBDriver derived using this high confidence list of mutations performed reasonably well across all five independent validation sets and produced 138 driver genes with sufficient literature evidence suggesting that our initial choice of the training dataset was overall beneficial. Second, since missense mutations are the most abundant form of somatic alterations [72], our machine learning models were all trained using missense mutations only. However, in principle, our approach could be extended to other types of mutations as well.

Additionally, during the external validation analysis, although NBDriver performed very well in terms of PPV (=0.941), the NPV (=0.608) was relatively low (Table 4A). To identify biologically relevant mutations for further functional validation, NPV is often overlooked as a classification metric. A high NPV allows us to exclude passenger mutations with greater confidence and reduces the number of driver mutations incorrectly labeled as passengers (false negatives). However, we observed that adding different combinations of multiple single predictors into ensemble models resulted in a significant improvement in the NPV (Table 4b). Our observations on the ensemble models’ performances were similar to those made by Martelotto *et al*. [38]. Last, we trained our machine learning models using the combined dataset containing mutational effects determined from experimental assays not specific to any cancer type. Hence, all our models were pan-cancer based. Consequently, a cancer-type specific analysis in the future would require the list of known driver and passenger mutations from specific tumor types.

## Conclusion

In this study, we showed that there is a significant difference in the nucleotide contexts surrounding driver and passenger mutations obtained from sequenced cancer genomes. Using efficient feature representation, we generated robust classification models that gave comparable performances across five independent validation sets. The predicted true positive mutations were part of genes with experimental support of being functionally relevant from multiple sources. Future experiments using a much larger sample size need to be performed to derive neighborhood-sequence-based classification scores for all possible missense mutations in the genome across several cancer types. This would be possible if future large-scale sequencing studies such as MSK-IMPACT [73]‘ PCAWG [44]‘ ICGC [7], and GENIE [74] produce a complete catalog of missense driver mutations with functional evidence in a cancer-type specific manner. This relatively novel strategy of utilizing the sequence neighborhoods for driver mutation identification can dramatically improve the annotation process’s efficiency for unknown mutations.

## Acknowledgements

This work was supported by Department of Biotechnology, Government of India (DBT) (BT/PR16710/BID/7/680/2016), IIT Madras, Initiative for Biological Systems Engineering (IBSE) and Robert Bosch Center for Data Science and Artificial Intelligence (RBC-DSAI).

